# Pleiotropic cellular responses underlying antibiotic tolerance in *Campylobacter jejuni*

**DOI:** 10.1101/2024.08.09.607412

**Authors:** Eunshin Cho, Jinshil Kim, Jeong In Hur, Sangryeol Ryu, Byeonghwa Jeon

## Abstract

Antibiotic tolerance enables antibiotic-susceptible bacteria to withstand prolonged exposure to high concentrations of antibiotics. Although antibiotic tolerance presents a major challenge for public health, its underlying molecular mechanisms remain to be elucidated. Previously, we have demonstrated that *Campylobacter jejuni* develops tolerance to clinically important antibiotics, including ciprofloxacin and tetracycline. To identify cellular responses associated with antibiotic tolerance, we conducted RNA-sequencing analysis on *C. jejuni* following the induction of antibiotic tolerance through exposure to elevated levels of ciprofloxacin or tetracycline. Additionally, we constructed knockout mutants for genes that showed significant changes in expression levels during antibiotic tolerance. We observed a significant upregulation of genes involved in protein chaperones, bacterial motility, DNA repair system, drug efflux pump, and iron homeostasis during antibiotic tolerance. Furthermore, the viability of these knockout mutants was significantly reduced compared to the wild-type strain, indicating the critical role of these cellular responses in sustaining antibiotic tolerance. Notably, we also found that the protein chaperone mutants Δ*dnaK*, Δ*clpB*, and Δ*groESL* exhibited increased protein aggregates under antibiotic treatment, suggesting protein chaperones play a critical role in managing protein aggregation and facilitating the survival of *C. jejuni* during antibiotic tolerance. In conclusion, our findings demonstrate that various cellular defense mechanisms collectively contribute to sustaining antibiotic tolerance in *C. jejuni*, providing novel insights into the molecular mechanisms of antibiotic tolerance that enable *C. jejuni* to withstand the lethal effects of antibiotics.

## INTRODUCTION

Antibiotic tolerance enables antibiotic-susceptible bacteria to survive antibiotic treatments without acquiring resistance, significantly compromising the effectiveness of antibiotic treatment (1, 2). Whereas antibiotic resistance involves genetic mutations and the acquisition of resistance genes, antibiotic tolerance is a transient state where bacteria can survive antibiotic exposure without genetic alternations or resistance acquisition (3, 4). Both antibiotic tolerance and persistence are common bacterial strategies to survive under antibiotic treatment and are often referred to interchangeably (3, 5). However, they represent distinct characteristics (5, 6).

Antibiotic persistence occurs in a small subpopulation of persister cells that are transiently tolerant to antibiotics and can resume growth upon removal of antibiotic treatment (5, 6). In contrast, antibiotic tolerance refers to a population-wide temporary state where bacteria can withstand and survive the toxic effects of antibiotics (3, 5, 7). Additionally, persistence is characterized by physiological dormancy as the lethal effects of antibiotics can be evaded in dormant bacteria with extremely slow metabolic and proliferation rates in response to antibiotics. In contrast, tolerance does not necessarily involve physiological dormancy (3, 5). Due to the difference, antibiotic persistence produces a unique biphasic time-kill curve pattern resulting from the survival of a tolerant subpopulation followed by the killing of the majority of non-tolerant bacterial populations, whereas tolerance generates a monophasic pattern that shows low levels of bacterial killing over time (3, 5, 7). Upon the removal of antibiotic stress, bacteria can be resuscitated to their normal physiological state. Recovery from antibiotic persistence from dormancy requires a relatively long time compared to that from tolerance (4, 8).

To investigate tolerance mechanisms using persister cells, specific procedures are required to selectively isolate and enrich a tolerant subpopulation (9–12). However, the transient nature and scarcity of persister cells present considerable challenges in researching antibiotic tolerance (13). Furthermore, most studies on tolerance have been constrained to a few hours of antibiotic treatment due to the rapid onset of bacterial death (14, 15), which may not accurately reflect clinical scenarios where pathogens are typically exposed to antibiotics over extended periods, ranging from days to weeks (16, 17). It is also critical to note that tolerance and persistence likely involve different cellular pathways, as they are distinct phenomena (3, 7). While persister cells survive antibiotics by entering a physiological state of dormancy and slowing metabolic processes, which may enable them to evade the lethal effects of antibiotics (18).

Furthermore, antibiotic tolerance can facilitate the development of antibiotic resistance as extended survival through tolerance can provide antibiotic-susceptible bacteria with the opportunity to acquire antibiotic resistance under antibiotic treatment (1, 2, 19). In a previous study, we discovered that *Campylobacter jejuni* develops tolerance when exposed to high concentrations of clinically important antibiotics, including ciprofloxacin (CIP) and tetracycline (TET) (20). *C. jejuni* is a leading bacterial cause of gastroenteritis, causing 400 million to 500 million infection cases worldwide per year (21). *Campylobacter* infections are generally self-limiting; however, antimicrobial therapies are required for severe infection cases, especially for the elderly and individuals with compromised immune systems (9–11). However, *C. jejuni* is increasingly resistant to clinically important antibiotics, particularly fluoroquinolones (FQs), the most commonly used oral antibiotic to treat various bacterial infections, including gastroenteritis (22, 23).

In our previous study, we demonstrated that high antibiotic concentrations promote the generation of reactive oxygen species (ROS) in *C. jejuni* during antibiotic tolerance, leading to DNA mutations resulting in antibiotic resistance, particularly FQ resistance (20). Moreover, we have found that antioxidation processes play a critical role in maintaining antibiotic tolerance in *C. jejuni*. Our current understanding of the molecular mechanisms of antibiotic tolerance is highly limited (8). Especially, there is a lack of information on how bacteria can address collateral cellular damage resulting from antibiotic treatment. *C. jejuni* offers a unique and feasible model for studying antibiotic tolerance due to its relatively faster growth compared to other tolerance-developing bacteria, such as *Mycobacterium tuberculosis* (24). Utilizing *C. jejuni*, in this study, we reveal the complex interplay of molecular processes that enable bacterial survival under high antibiotic concentrations through tolerance.

## RESULTS

### Transcriptome changes during antibiotic tolerance in *C. jejuni*

In order to understand transcriptome changes underlying antibiotic tolerance, we performed RNA-Seq after exposing *C. jejuni* to high concentrations of CIP or TET. These two antibiotics were chosen based on our previous findings that *C. jejuni* can survive in the presence of high concentrations of these antibiotics through tolerance for an extended period (20). Notably, CIP and TET have different modes of action, with CIP disrupting bacterial DNA synthesis by targeting DNA gyrase (25) and TET inhibiting protein synthesis by targeting the 30S subunit of ribosomes (26). Since antimicrobial processes by an antibiotic can cause transcriptomic changes, using the two tolerance-inducing antibiotics with different modes of action will contribute to the identification of cellular responses unique to antibiotic tolerance. Upon exposure to 100x MICs of CIP, *C. jejuni* underwent viability reduction for 2 days, followed by the emergence of CIP-resistant populations (Fig. 1A). The pattern of *C. jejuni* survival in the presence of these antibiotics is different from that observed in *Escherichia coli*, where CIP significantly reduced viability within a few hours, followed by the survival of persister cells at extremely low levels (Fig. 1B).

**Fig. 1.**
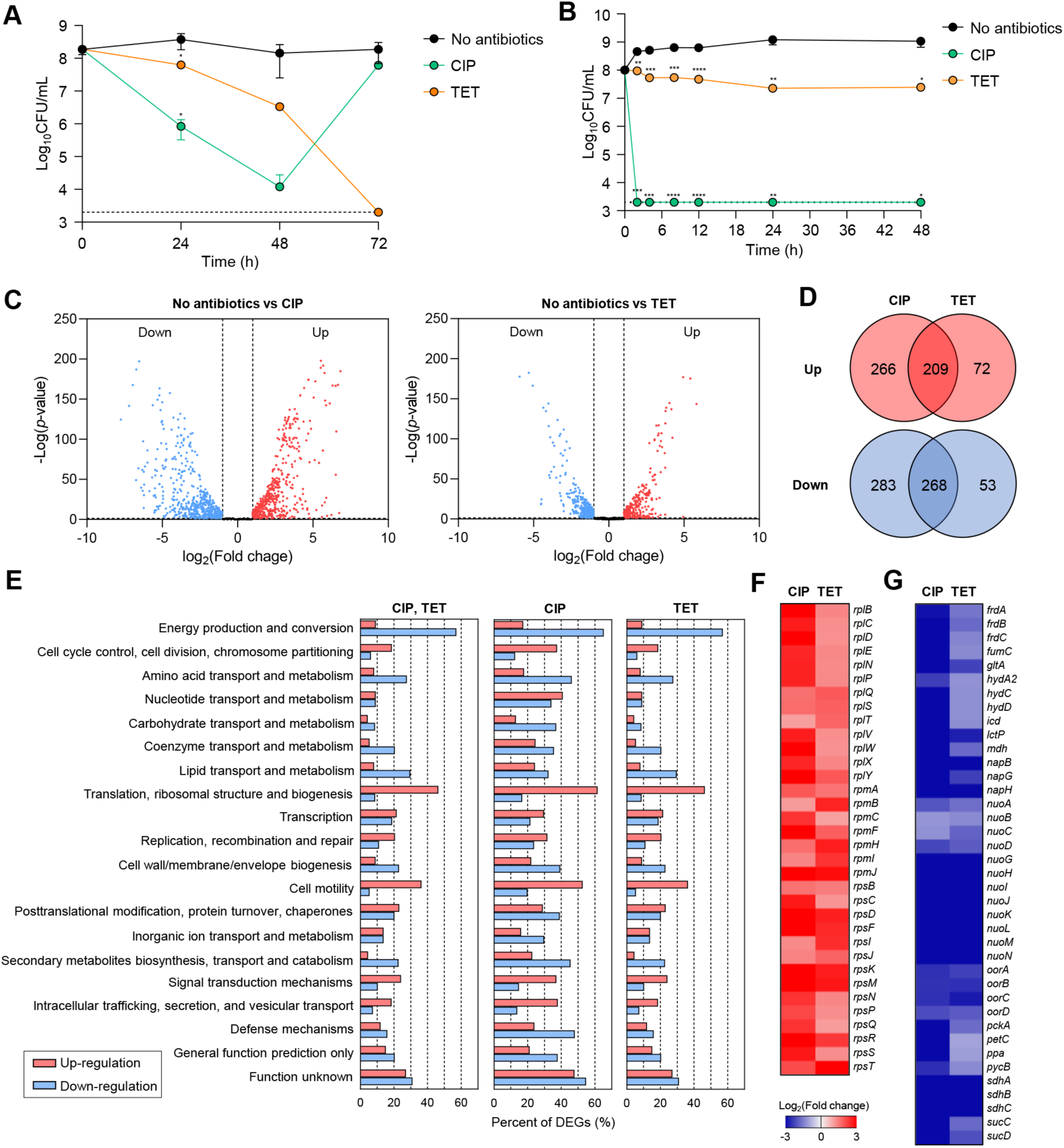
Antibiotic tolerance and transcriptomic changes in *C. jejuni* in the presence of high concentrations of ciprofloxacin (CIP) or tetracycline (TET). (A-B) Induction of antibiotic tolerance in *C. jejuni* (A) and *E. coli* (B) by exposure to 100x MICs of CIP (6.3 μg/ml and 1.6 μg/ml, respectively) or TET (3.1 μg/ml and 50 μg/ml, respectively). Error bars represent the standard deviations of three independent experiments. The data were statistically analyzed by the Student’s *t* test in comparison with the untreated control (No antibiotics); *, *P* < 0.05; **, *P* < 0.01; ***, *P* < 0.001; ****, *P* < 0.0001. (C-G) The transcriptomic changes in *C. jejuni* after exposure to 100x MICs of CIP (6.3 μg/ml) or TET (3.1 μg/ml) based on RNA-Seq. Fold change was defined compared to the untreated control. Volcano plots (C) and Venn diagrams (D) depicting differentially expressed genes (DEGs) in *C. jejuni* after exposure to 100x MICs of CIP or TET. The percentages of DEGs (E) in *C. jejuni* during antibiotic tolerance. Heat maps of the genes associated with translation, ribosomal structure, and biogenesis (F) or energy production and conversion (G) after exposure to 100x MICs of CIP or TET. The heat maps were constructed with Gitools.

Moreover, high TET concentrations led to killing of *C. jejuni* (Fig. 1A), whereas TET exhibited only bacteriostatic activity in *E. coli* (Fig. 1B). The results show that *C. jejuni* is significantly tolerant to TET and CIP.

The results of the RNA-Seq analysis have revealed that significant transcriptome changes occur during antibiotic tolerance. Compared to the untreated sample, CIP was found to induce differential expression in a total of 1,026 genes, including 475 upregulated genes and 551 downregulated genes (Fig. 1C-D), whereas TET exhibited differential expression in a total of 602 genes, with 281 upregulated genes and 321 downregulated genes (Fig. 1C-D). Additionally, a total of 477 genes were differentially expressed in both CIP and TET (Fig. 1D). Notably, CIP upregulated genes involved in various cellular functions, such as cell cycle control, cell division, chromosome partitioning, and cell motility, while downregulating genes associated with amino acid transport and metabolism, secondary metabolites biosynthesis, and defense mechanisms (Fig. 1E). On the other hand, both CIP and TET commonly increased the transcription of genes associated with translation and ribosomal structure, while downregulating those related to energy production and conversion (Fig. 1E-G). Notably, genes involved in protein translation and tRNA genes were upregulated (Fig. 1F), indicating that *C. jejuni* is not dormant during antibiotic tolerance and actively synthesizes proteins to respond to antibiotic stress. The increased expression of these genes may contribute to antibiotic tolerance in *C. jejuni*.

### Protein chaperones contribute to sustaining antibiotic tolerance

Protein chaperones are critical for assisting in proper protein folding and disaggregation, particularly under stressful conditions (27). The three major bacterial chaperone complexes are the trigger factor, the DnaK-DnaJ, and the GroEL-GroES complexes (28). RNA-Seq analysis has revealed that genes encoding these chaperone complexes, specifically *clpB*, *dnaK*, *groES*, *groEL*, and *tig*, are significantly upregulated during antibiotic tolerance (Fig. 2A), suggesting that these chaperones play an important role in bacterial survival under antibiotic treatment. Notably, Δ*dnaK*, Δ*clpB*, and Δ*groESL* mutations have been shown to compromise antibiotic tolerance, resulting in significant viability reductions after 48 h of antibiotic exposure (Fig. 2B). This was particularly evident under treatment with TET, where Δ*clpB* and Δ*groESL* mutations facilitated bacterial cell death (Fig. 2B).

**Fig. 2.**
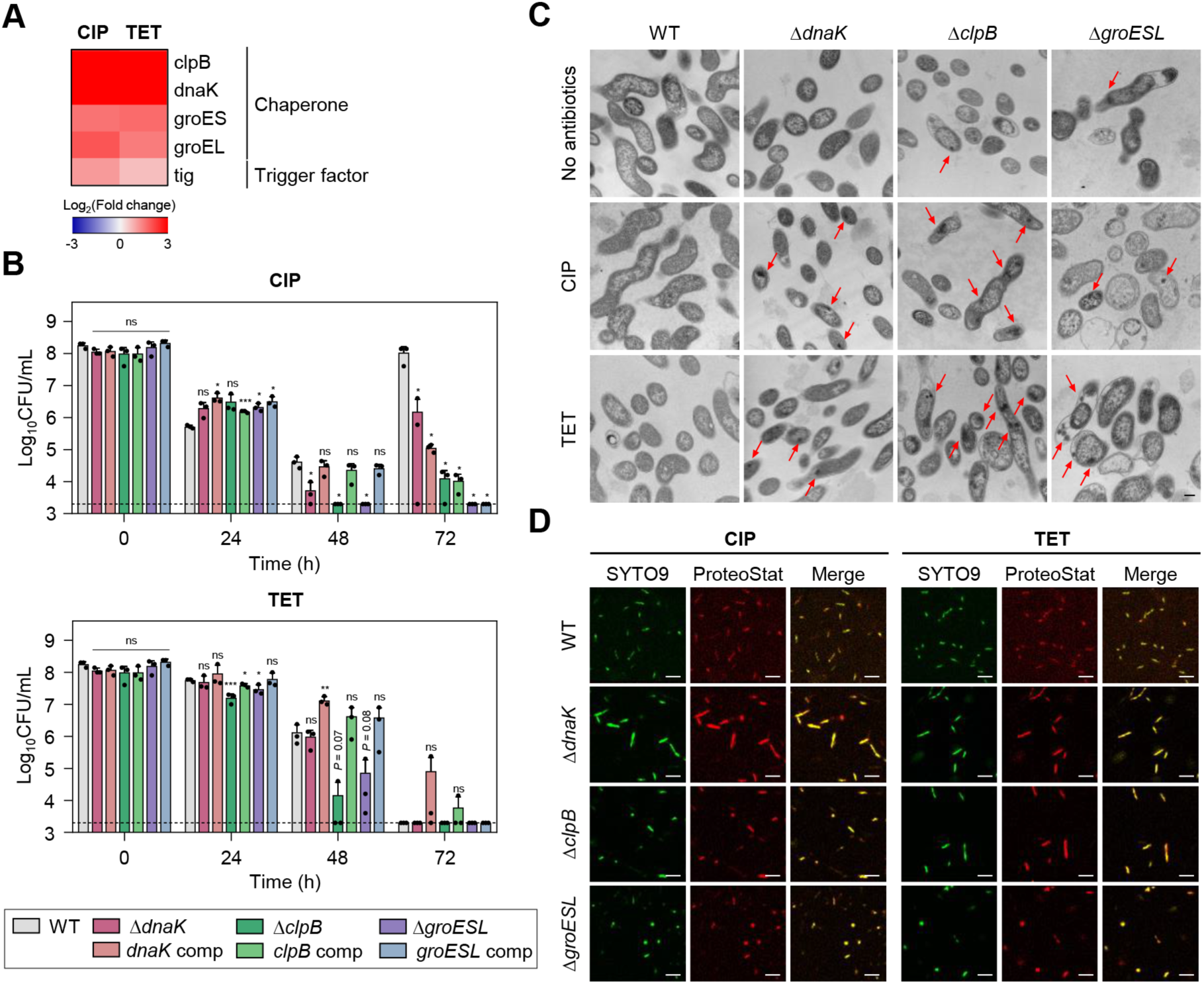
Effects of chaperones on antibiotic tolerance in *C. jejuni* in the presence of high concentrations of ciprofloxacin (CIP) or tetracycline (TET). (A) Heat maps of the genes associated with chaperones after exposure to 100x MICs of CIP (6.3 μg/ml) or TET (3.1 μg/ml). The heat maps were constructed with Gitools. (B) Induction of antibiotic tolerance by exposure to 100x MICs of CIP (6.3 μg/ml) or TET (3.1 μg/ml). Error bars represent the standard deviations of three independent experiments. The data were statistically analyzed by the Student’s *t* test in comparison with WT. *, *P* < 0.05; **, *P* < 0.01; ***, *P* < 0.001; ns, not significant; *dnaK* comp, *dnaK*-complemented strain; *clpB* comp, *clpB*-complemented strain; *groESL* comp, *groESL*- complemented strain. (C) Formation of protein aggregates induced after exposure to 100x MICs of CIP or TET. Red arrows indicated protein aggregates observed by cross-section transmission electron microscopy (TEM). The formation of protein aggregates was compared to the untreated control (No antibiotics). The scale bar represents 0.2 μm. (D) Protein aggregates were visualized with fluorescent probes. Live cells were stained with SYTO9 (green) and protein aggregates with ProteoStat (red). The merged images are shown in yellow. The scale bar represents 5 μm.

Environmental stress can disrupt protein homeostasis, leading to the formation of insoluble protein aggregates within bacterial cells (29, 30). Transmission electron microscopy (TEM) has been employed to investigate whether antibiotic exposure induces protein aggregation during antibiotic tolerance. Compared to wild type (WT), more protein aggregates were detected in Δ*dnaK*, Δ*clpB*, and Δ*groESL* mutants (Fig. 2C). Additionally, the use of ProteoStat, a fluorescent dye that selectively binds to misfolded and aggregated proteins (31), has corroborated these observations. Consistently, enhanced protein aggregation (red in Fig. 2D) was observed in the chaperone mutants compared to WT. In contrast, there was no discernible difference in negative controls without antibiotic treatment (Fig. S3). These findings indicate the critical function of chaperone proteins in bacterial survival during antibiotic tolerance.

### Association of bacterial motility with antibiotic tolerance

The motility of *C. jejuni* is facilitated by polar flagella that consist of flagellins encoded by *flaA* and *flaB*, which are regulated by the sigma factors FliA and RpoN, respectively (32, 33). Among these flagellin genes, *flaA* is the major flagellin as its inactivation leads to a loss of motility and a decrease in virulence, whereas an inactivation of *flaB* results in the formation of truncated flagella but does not reduce motility (33). Genes related to motility were significantly upregulated during tolerance (Fig. 3A), suggesting that bacterial motility is critical for antibiotic tolerance.

**Fig. 3.**
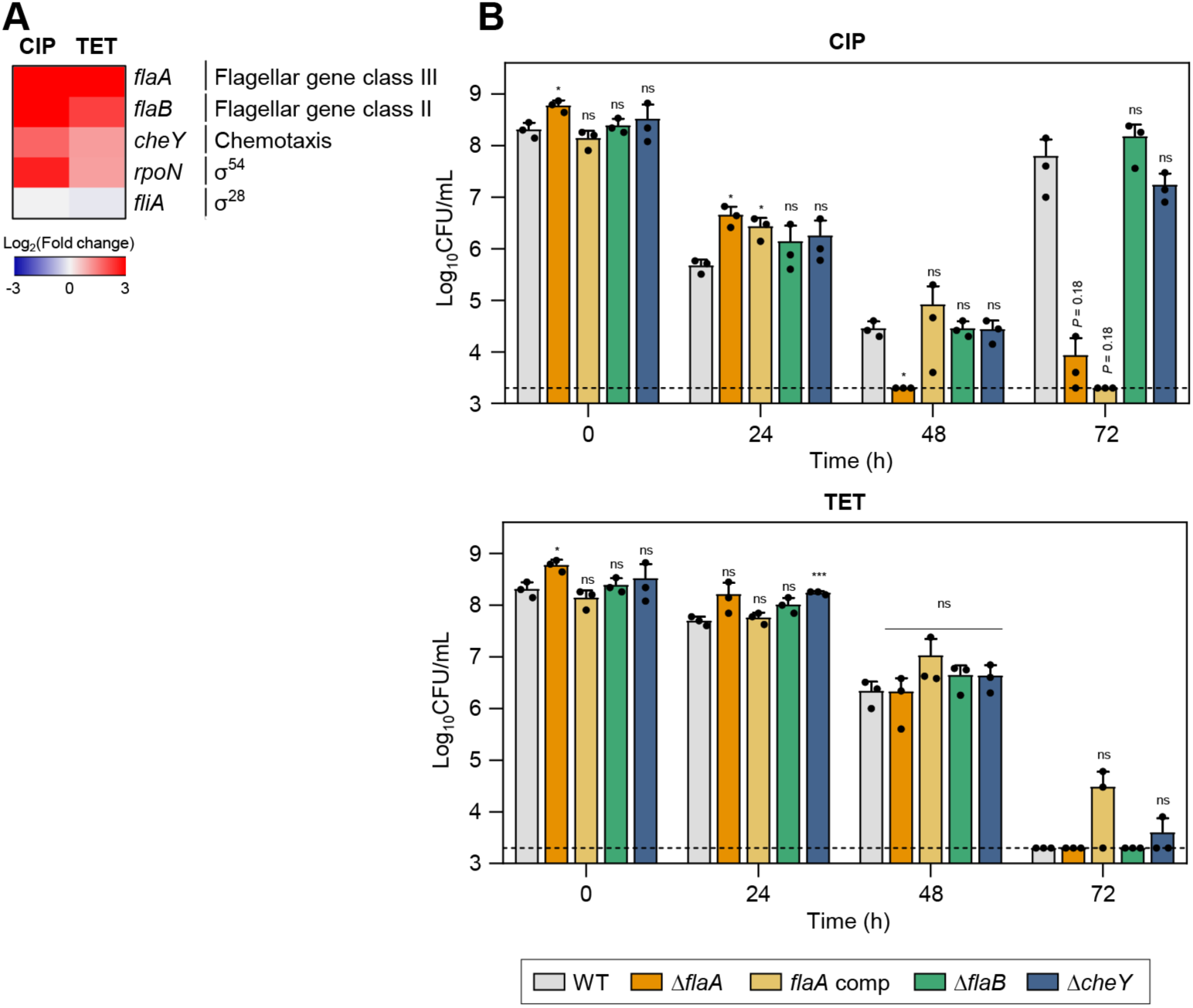
Effects of motility-related genes on antibiotic tolerance in *C. jejuni* in the presence of high concentrations of ciprofloxacin (CIP) or tetracycline (TET). (A) Heat maps of the genes associated with motility after exposure to 100x MICs of CIP (6.3 μg/ml) or TET (3.1 μg/ml). The heat maps were constructed with Gitools. (B) Induction of antibiotic tolerance by exposure to 100x MICs of CIP (6.3 μg/ml) or TET (3.1 μg/ml). Error bars represent the standard deviations of three independent experiments. The data were statistically analyzed by the Student’s *t* test in comparison with WT; *, *P* < 0.05; ***, *P* < 0.001; ns, not significant; *flaA* comp, *flaA*-complemented strain.

Additionally, the *cheY* gene, which is integral to bacterial chemotaxis (34), was also upregulated during antibiotic tolerance (Fig. 3A).

To better understand the contributions of motility and chemotaxis to antibiotic tolerance, we constructed Δ*flaA*, Δ*flaB*, and Δ*cheY* mutants. When treated with CIP, the Δ*flaA* mutant demonstrated a significant reduction in viability compared to WT, suggesting that *flaA*-mediated motility is crucial for CIP tolerance. Under TET treatment, there was no observable difference in viability reduction between the mutants and WT (Fig. 3B). This indicates that the role of motility and chemotaxis in antibiotic tolerance may vary depending on the antibiotic.

### DNA repairs are critical for antibiotic tolerance

A number of genes of DNA repairs were significantly upregulated during antibiotic tolerance (Fig. 4A). Specifically, the *ssb* gene, which encodes the single-stranded DNA-binding protein crucial for DNA replication, recombination, and repair (35), was upregulated by exposure to high concentrations of antibiotics. Additionally, antibiotic treatment also upregulated *dprA* encoding DNA processing protein A (DprA), which assists in the integration of single-stranded DNA into the genome and is involved in natural transformation in *C. jejuni* (36). Interestingly, antibiotic treatment did not upregulate *recA* (Fig. 4A), presumably due to the absence of SOS response systems in *C. jejuni* (37).

**Fig. 4.**
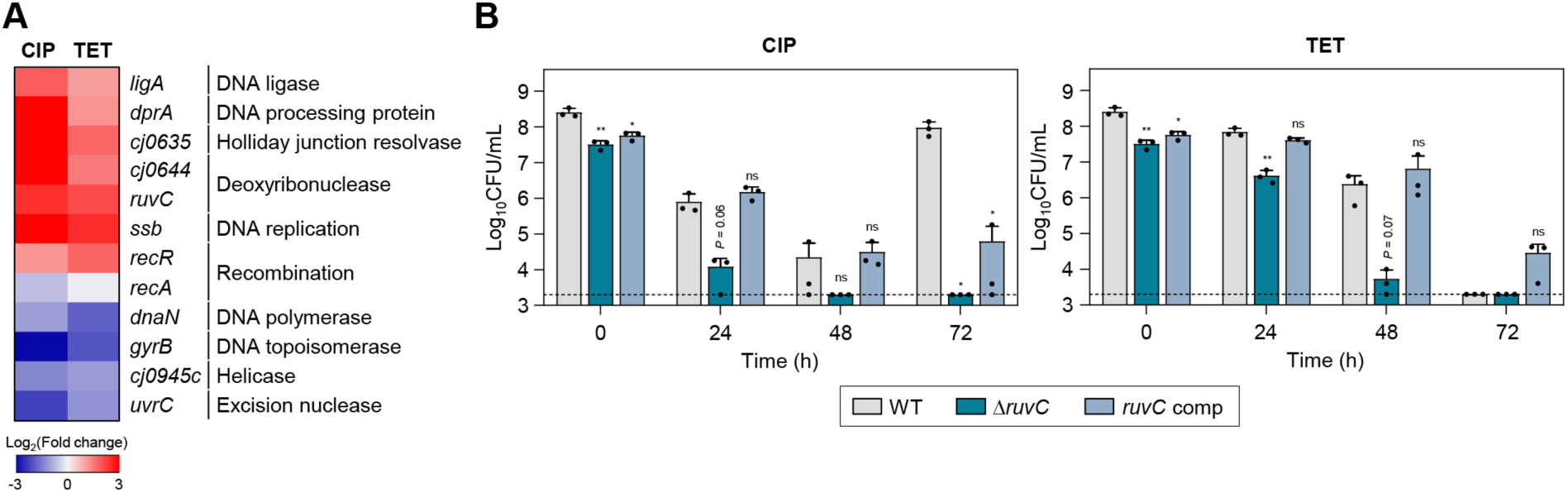
Effects of DNA repair genes on antibiotic tolerance in *C. jejuni* in the presence of high concentrations of ciprofloxacin (CIP) or tetracycline (TET). (A) Heat maps of the genes associated with DNA repair after exposure to 100x MICs of CIP (6.3 μg/ml) or TET (3.1 μg/ml). The heat maps were constructed with Gitools. (B) Induction of antibiotic tolerance by exposure to 100x MICs of CIP (6.3 μg/ml) or TET (3.1 μg/ml). Error bars represent the standard deviations of three independent experiments. The data were statistically analyzed by the Student’s *t* test in comparison with WT; *, *P* < 0.05; **, *P* < 0.01; ns, not significant; *ruvC* comp, *ruvC*-complemented strain.

To evaluate the impact of DNA repair systems on antibiotic tolerance, we constructed a Δ*ruvC* mutant and measured its viability in the presence of 100x MICs of CIP or TET. RuvC plays an important role in DNA repair and recombination by cleaving Holliday junctions (38). The *ruvC* was selected because its transcriptional level was significantly enhanced by both CIP and TET (Fig. 4A). Moreover, the Δ*ruvC* mutation severely impaired antibiotic tolerance in *C. jejuni* (Fig. 4B). Additionally, the Δ*ruvC* mutation sensitized *C. jejuni* to antibiotics, displaying a decrease in the MICs of CIP and TET by 4-fold and 2-fold, respectively (Table S1).

It is noteworthy that the upregulation of DNA repair genes was observed during tolerance induced by TET, an antibiotic that targets protein synthesis (Fig. 4A). Furthermore, the inactivation of *ruvC* markedly diminished bacterial viability in the presence of TET. These findings indicate the importance of DNA repair processes in maintaining antibiotic tolerance, regardless of the mode of action of an antibiotic used for tolerance induction.

### Drug efflux pumps contribute to maintaining antibiotic tolerance

Drug efflux pumps play a critical role in antibiotic resistance by reducing the intracellular concentration of antibiotics (39). In *C. jejuni*, CmeABC is the primary efflux system that confers resistance across various antibiotic classes (40). CmeDEF is another drug efflux pump that operates alongside CmeABC to maintain cell viability under antibiotic treatment (41). Transcriptomic analysis showed that the expression of these efflux pump genes is modulated in response to antibiotic exposure (Fig. S2). To further elucidate the role of these pumps during antibiotic tolerance, we constructed Δ*cmeC* and Δ*cmeF* mutants. CmeABC and CmeDEF are the resistance-nodulation-cell division (RND)- type efflux pumps, which are composed of three proteins spanning the cytoplasmic space and both cell membranes; thus, the absence of any one component renders the entire pump nonfunctional. Under CIP treatment, the viability reductions in these efflux pump mutants were slightly more substantial compared to WT after 48 h (Fig. 5A). In contrast to WT, notably, the emergence of FQ-resistant strains was not observed in the Δ*cmeC* mutant after 72 h (Fig. 5A). In the presence of 100x MICs of TET, tolerance was significantly compromised in both Δ*cmeC* and Δ*cmeF* mutants (Fig. 5A).

**Fig. 5.**
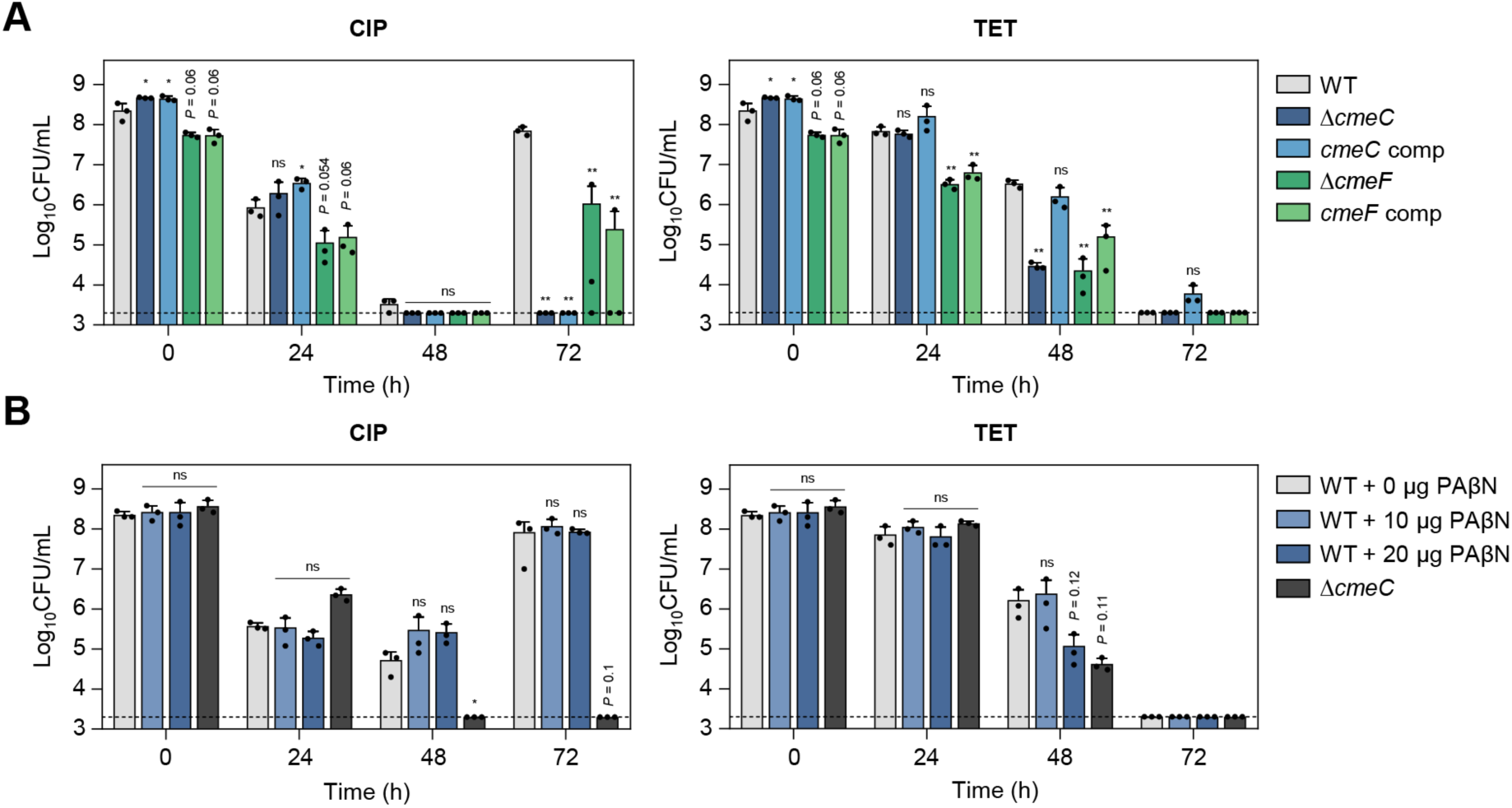
Effects of drug efflux pumps on antibiotic tolerance in *C. jejuni* in the presence of high concentrations of ciprofloxacin (CIP) or tetracycline (TET). (A-B) Induction of antibiotic tolerance in efflux pump knockout mutants (A) by exposure to 100x MICs of CIP (6.3 μg/ml) or TET (3.1 μg/ml). Induction of antibiotic tolerance in WT in the presence of phenylalanine-arginine β-naphthylamide (PAβN), an efflux pump inhibitor (B) after exposure to 100x MICs of CIP (6.3 μg/ml) or TET (3.1 μg/ml). Error bars represent the standard deviations of three independent experiments. The data were statistically analyzed by the Student’s *t* test in comparison with WT; *, *P* < 0.05; **, *P* < 0.01; ns, not significant; *cmeC* comp, *cmeC*-complemented strain; *cmeF* comp, *cmeF*-complemented strain.

The role of efflux pumps in antibiotic tolerance was also assessed using the efflux pump inhibitor phenylalanine-arginine β-naphthylamide (PAβN). Similarly, PAβN did not affect bacterial viability during CIP-induced tolerance, whereas PAβN significantly reduced viability in the presence of 100x MICs of TET (Fig. 5B). These results suggest that drug efflux pumps may contribute to antibiotic tolerance depending on the antibiotic.

### Increased iron accumulation during antibiotic tolerance

RNA-Seq analysis has also revealed a notable upregulation of *fur* transcription during antibiotic tolerance, particularly when tolerance was induced by CIP (Fig. 6A). A Δ*fur* mutation significantly compromised the viability in the presence of 100x MICs of TET and reduced the emergence of FQ-resistant strains under the treatment with 100x MICs of CIP (Fig. 6B). Interestingly, the level of intracellular iron is significantly increased during antibiotic tolerance (Fig. 6C). Iron is a cofactor of a range of proteins essential for fundamental physiological processes (42, 43). Most Gram-negative bacteria, including *C. jejuni*, maintain cytoplasmic iron levels using Fur, a transcriptional repressor (44, 45). Iron exists in either the reduced ferrous form (Fe^2+^) or the oxidized ferric form (Fe^3+^). Fe^2+^ can passively diffuse through the outer-membrane porins and is imported by FeoB, which is the only Fe^2+^ transport system that has been identified in *C. jejuni* (46). Fe^3+^ is imported through specific ligand-gated outer-membrane receptor proteins using siderophores, including Fe^3+^-enterochelin (CeuBCDE, CfrA) and Fe^3+^-rhodotorulic acid (p19, Cj1658-63) (47, 48), During antibiotic tolerance, genes for Fe^3+^-uptake systems involving rhodotorulic acid and hemin, and the Fe^2+^-uptake FeoB were also down-regulated (Fig. 6A). Presumably, *fur* transcription was increased so that Fur can prevent further iron uptake to maintain iron homeostasis and reduce cellular toxicity. After acquisition, iron can be stored in the form of ferritin or incorporated into iron-sulfur complexes (49, 50). Moreover, the transcription of *dps*, involved in the sequestration of intracellular free iron (51), is significantly increased during tolerance, which is aligned with cellular changes to mitigate cellular toxicity caused by increases in iron levels. These findings suggest that during antibiotic tolerance, *C. jejuni* actively modulates its iron acquisition and storage systems to mitigate the deleterious effects of iron overload. The coordinated regulation of iron uptake and storage genes reflects *C. jejuni*’s adaptive response to maintain iron homeostasis during antibiotic tolerance.

**Fig. 6.**
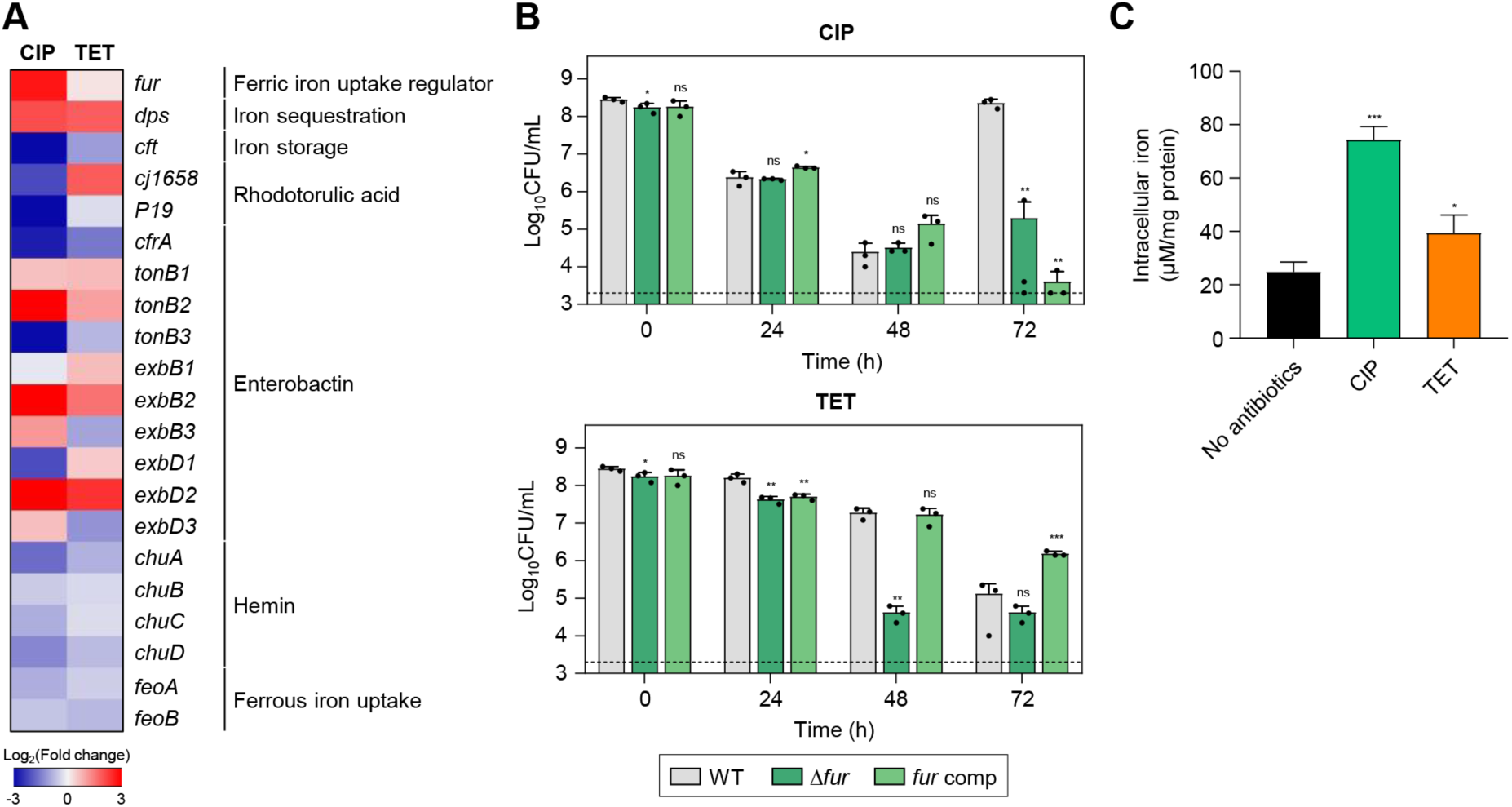
Effects of iron accumulation in *C. jejuni* in the presence of high concentrations of ciprofloxacin (CIP) or tetracycline (TET). (A) Heat maps of the genes associated with iron metabolism after exposure to 100x MICs of CIP (6.3 μg/ml) or TET (3.1 μg/ml). The heat maps were constructed with Gitools. (B) Induction of antibiotic tolerance by exposure to 100x MICs of CIP (6.3 μg/ml) or TET (3.1 μg/ml). Error bars represent the standard deviations of three independent experiments. The data were statistically analyzed by the Student’s *t* test in comparison with WT; *, *P* < 0.05; **, *P* < 0.01; ***, *P* < 0.001; ns, not significant; *fur* comp, *fur*-complemented strain. (C) The intracellular iron content in *C. jejuni* after exposure to 100x MICs of CIP (6.3 μg/ml) or TET (3.1 μg/ml). Error bars represent the standard deviations of three independent experiments. The data were statistically analyzed by the Student’s *t* test in comparison with the untreated control (No antibiotics); *, *P* < 0.05; ***, *P* < 0.001.

## DISCUSSION

Here, we provide an extensive analysis of the cellular mechanisms that contribute to *C. jejuni*’s ability to tolerate high antibiotic concentrations, revealing that antibiotic tolerance involves various defense mechanisms, including protein chaperones, bacterial motility, DNA repair systems, drug efflux pumps, and iron homeostasis. Notably, we discovered that the function of protein chaperones is critical for antibiotic tolerance. Under environmental stress, bacteria may experience protein aggregation, which perturbs protein homeostasis and leads to the formation of insoluble protein aggregates (29, 30). Although protein aggregation is generally associated with negative cellular outcomes, such as impaired cellular functions and cell death, it is also considered a survival strategy against antibiotic treatment by triggering bacterial dormancy in persister cells (18). When antibiotics are removed, persister cells must resolve protein aggregates to revive and resume growth, which is a process that determines the lag time for bacterial regrowth (52). Similarly, persister-enriched *Staphylococcus aureus* populations contain increased levels of insoluble protein aggregates (53). These findings indicate a correlation between protein aggregation and bacterial dormancy under antibiotic stress.

Persister cells use dormancy to evade the lethal effects of antibiotics due to the extremely slow metabolic processes (18). In the state of antibiotic persistence, DnaK and ClpB contribute to maintaining a dormant state, allowing persister cells to survive antibiotic challenges and later resuscitate into actively growing bacteria upon the removal of antibiotic stress (52). DnaK is a chaperone protein that recognizes and binds to exposed hydrophobic regions on partially misfolded proteins (54, 55). It then transfers these partially folded proteins to ClpB, an AAA+ ATPase chaperone, which disaggregates and solubilizes aggregated proteins (56). The ClpB/DnaK bi-chaperone system interplays to address protein misfolding and aggregation issues, preventing the formation of toxic protein aggregates in response to stress conditions, including antibiotic treatment (57, 58). While the role of protein aggregation in inducing physiological dormancy and the function of chaperone proteins in antibiotic resistance has been documented (52), this mechanism may not be applied to understanding antibiotic tolerance. Tolerance is characterized by a uniform response throughout a bacterial population that does not necessarily involve dormancy, as opposed to persistence which is specific to a dormant subpopulation (3, 5). Our findings suggest that chaperone proteins should disaggregate protein aggregates to maintain critical cellular functions for bacterial viability during antibiotic tolerance. Although the precise role of molecular chaperones during antibiotic tolerance remains unexplained, our data suggest that protein chaperones contribute to bacterial survival under antibiotic treatment by contributing to protein disaggregation through tolerance.

We conducted transcriptome analysis after antibiotic exposure for 24 h to capture the effects of extended antibiotic exposure on gene expression. Notably, the transcriptional levels of chaperon genes were increased over the duration of antibiotic exposure from 2 h to 24 h (Fig. S4), indicating the critical function of chaperons in maintaining antibiotic tolerance. Moreover, chaperone mutations did not reduce viability at 24 h but rendered the mutants significantly vulnerable to antibiotics after 48 h. This highlights the importance of protein chaperones in antibiotic tolerance and consequently bacterial survival during prolonged antibiotic exposure.

Our findings further demonstrate the pivotal role of DNA repair in sustaining antibiotic tolerance. Within bacterial populations enriched with persister cells of *S. aureus*, DNA repair proteins are downregulated, which indicates physiological dormancy in persister cells (53). In contrast, we observed the upregulation of a range of DNA repair genes, which implies that *C. jejuni* is not dormant during tolerance and should preserve DNA integrity, potentially to support vital physiological functions. It is noteworthy that *C. jejuni* does not have the SOS response system which is critical for addressing DNA damage in various bacteria (37). Instead, *C. jejuni* seems to possess alternative DNA repair mechanisms that are independent of the SOS response (59). During antibiotic tolerance, high antibiotic concentrations lead to the generation of toxic ROS, particularly hydroxyl radicals, which can damage DNA (20). Presumably, DNA repair contributes to addressing DNA damage and mutations, especially those that interrupt the function of essential genes, thereby ensuring the maintenance of bacterial survival.

Notably, our research has uncovered that the concentration of intracellular iron is increased during antibiotic tolerance (Fig. 6C). Although the mechanisms behind the increase in iron levels during tolerance have yet to be elucidated, the interaction of intracellular free iron with hydrogen peroxide leads to the production of DNA-damaging hydroxyl radicals via the Fenton reaction (60). To mitigate oxidative stress associated with increased intracellular iron levels during antibiotic tolerance, *C. jejuni* should prevent further iron uptake and sequester intracellular free iron. To this end, we observed that the transcriptional levels of *fur* (a Fe^3+^ uptake repressor) and *dps* (an intracellular iron sequestration protein) were markedly increased, whereas *feoB* encoding a ferrous iron transporter was down-regulated (Fig. 6A). Our previous study also has demonstrated that oxidative stress is highly elevated in *C. jejuni* during tolerance and that antioxidant enzymes, such as alkyl hydroperoxide reductase (AhpC), play a critical role in maintaining antibiotic tolerance (20). The critical function of iron in the generation of hydroxyl radicals indicates that maintaining iron homeostasis is essential for sustaining antibiotic tolerance, presumably through the control of oxidative stress.

In summary, our study first revealed pleiotropic cellular responses that drive bacterial tolerance to antibiotics in *C. jejuni*. Our findings demonstrate that *C. jejuni* extensively utilizes various cellular defense mechanisms, including antioxidation, protein chaperoning, DNA repair pathways, drug efflux pumps, and iron homeostasis to survive antibiotic treatment (Fig. 7).

**Fig. 7.**
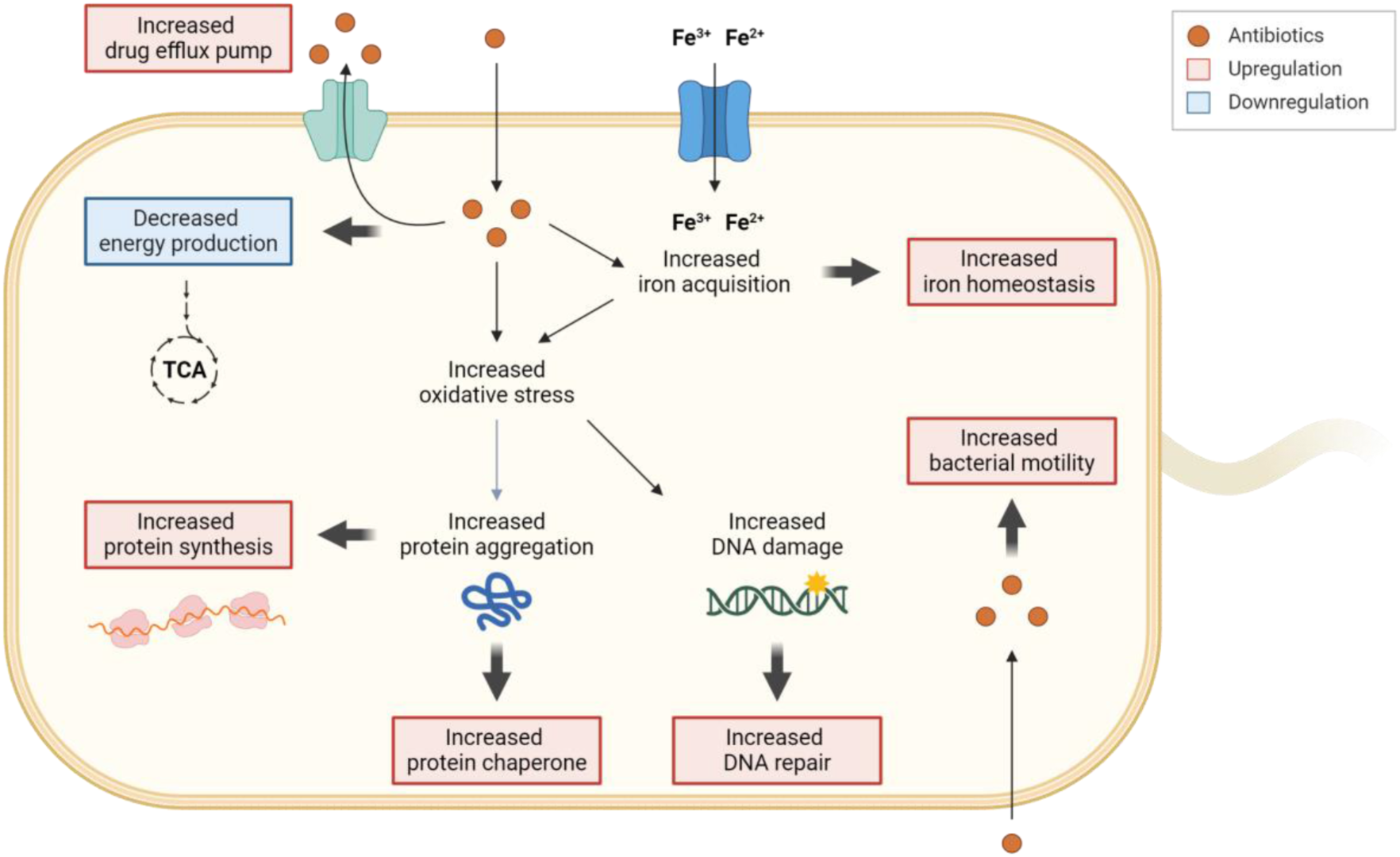
Schematic representation of the cellular responses involved in the development of antibiotic tolerance in *C. jejuni*. When exposed to high levels of antibiotics, drug efflux pumps actively work to decrease the intracellular concentrations of antibiotics. Exposure to high concentrations of antibiotics leads to an accumulation of iron within the cell, subsequently influencing the gene expression of iron acquisition and storage systems to maintain iron homeostasis. The increased oxidative stress caused by antibiotics induces DNA damage, prompting the upregulation of the DNA repair system. Furthermore, oxidative stress may also trigger protein aggregation, resulting in the upregulation of protein synthesis and protein chaperones to disaggregate protein aggregates. Intriguingly, intracellular antibiotics may also stimulate bacterial motility. Additionally, exposure to high concentrations of antibiotics leads to the downregulation of genes associated with energy production. The transcriptional changes are visually represented with bold arrows, with upregulation highlighted in red and downregulation in blue.

Notably, iron homeostasis can be associated with antibiotic tolerance by modulating intracellular iron levels. Future studies await to evaluate the effects of intracellular free iron on tolerance maintenance and to further clarify the underlying molecular mechanisms.

## MATERIALS AND METHODS

### Bacterial strains and growth conditions

*C. jejuni* NCTC 11168 was used as WT in this study. *C. jejuni* strains were grown microaerobically (5% O2, 10% CO2, and 85% N2) at 42°C on Mueller-Hinton (MH) media (Oxoid, Hampshire, UK). *E. coli* strains were grown at 37°C on Luria-Bertani (LB) media (Difco, MI, USA). Occasionally, culture media were supplemented with antibiotics, including carbenicillin (100 μg/ml), kanamycin (50 μg/ml), and chloramphenicol (12.5 μg/ml).

### Time-kill assay

Overnight cultures of *C. jejuni* grown on MH agar were resuspended in 5 ml of MH broth in a 14-ml round-bottom tube (BD Falcon, MA, USA) to an optical density at 600 nm (OD600) of 0.08. The bacterial suspension was then incubated with shaking under microaerobic conditions. After 7 h incubation, antibiotic exposure was initiated by adding 100x MICs of CIP (6.3 μg/ml) or TET (3.1 μg/ml). After 24, 48, and 72 h incubation, 100 μl of *C. jejuni* cultures were harvested and washed with ice-cold phosphate-buffered saline (PBS) three times. After washing, bacterial cells were resuspended in 1 ml of PBS and diluted with MH broth. Five microliters of bacterial cells were spotted onto MH agar and incubated for two days to assess viability. To examine the effect of an efflux pump inhibitor on antibiotic tolerance, *C. jejuni* was incubated with PAβN (10 or 20 μg/ml) for 7 h in the presence of antibiotics as described above. The assay was also conducted with *E. coli*. *E. coli* MG1655 was incubated in LB broth with shaking under aerobic conditions. At the exponential growth phase, the bacterial population was adjusted to a concentration of 10^8^ CFU/ml. Subsequently, the cultures were treated with 100x MICs of CIP (1.6 μg/ml) or TET (50 μg/ml). The samples were harvested after incubation for 2, 4, 8, 12, 24, and 48 h. After washing with ice-cold PBS three times, the bacterial cells were resuspended in PBS, diluted, and spotted onto the LB agar plates. After 12 h incubation, bacterial viability was assessed.

### Total RNA extraction, RNA-seq, and analysis

Overnight cultures of *C. jejuni* NCTC 11168 grown on MH agar were harvested and suspended in MH broth to an OD600 of 0.08. A 3 ml bacterial suspension in a 19 ml glass culture tube was incubated for 7 h with shaking under microaerobic conditions. After 7 h, cultures were treated with 100x MICs of either CIP (6.3 μg/ml) or TET (3.1 μg/ml) for 24 h. Bacterial cultures (2.5 ml) were treated with 5% ice-cold phenol-ethanol solution, and total bacterial RNAs were isolated using the RNeasy Minikit (Qiagen, Hilden, Germany) according to the manufacturer’s instructions. The quantity and quality of total RNA samples were assessed using a NanoPhotometer N60 (Implen, Munich, Germany), and three biological replicate RNA samples were sent to Macrogen (Seoul, Republic of Korea) for RNA sequencing.

The quality and quantity of total RNA were further evaluated using an Agilent Technologies 2100 Bioanalyzer, ensuring a RNA integrity number (RIN) value greater than 7. A library was prepared independently with 1 μg of total RNA for each sample by Illumina TruSeq Stranded mRNA Sample Prep Kit (Illumina, Inc., CA, USA). Initially, bacterial rRNA-depleted samples were prepared by using the NEBNext rRNA Depletion kit (NEB, NA, USA). After rRNA depletion, the remaining RNA was fragmented into small pieces using divalent cations under elevated temperature. The RNA fragments were converted into first-strand cDNA using SuperScript II reverse transcriptase (Invitrogen, MA, USA) and random primers. This is followed by second-strand cDNA synthesis using DNA Polymerase I, RNase H and dUTP. These cDNA fragments underwent an end repair process, the addition of a single ‘A’ base, and ligation of the adapters. The products were then purified, enriched with PCR, and processed to create the final cDNA library. Library quantification was carried out using KAPA Library Quantification kits for Illumina Sequencing platforms, and qualification was performed using the TapeStation D1000 ScreenTape (Agilent, CA, USA). Indexed libraries were then submitted to an Illumina NovaSeq 6000 (Illumina, Inc., CA, USA), employing paired-end (2×100 bp) sequencing by Macrogen (Seoul, Republic of Korea).

The expression level of each gene was normalized by calculating reads per kilobase per million mapped reads (RPKM) using CLC Workbench. Fold change was determined in comparison to the untreated control (No antibiotics). Differentially expressed genes (DEGs; fold change≥ 2 or ≤-2; *P* < 0.05) were filtered and visualized using the Gitools.

### Quantitative real-time PCR (qRT-PCR)

Total RNA was extracted as described above, and cDNA was synthesized using cDNA EcoDry premix (Clontech, USA). qRT-PCR was performed in a 20 μl reaction volume containing cDNA, iQ SYBR Green supermix (Bio-Rad, USA), and each primer, using the CFX Connect real-time PCR detection system (Bio-Rad, USA). The primer sets used in qRT-PCR are described in Table S2. The cycling conditions were as follows: 95°C for 5 min; 40 cycles at 95°C for 15 s, 55°C for 15 s, and 72°C for 30 s; followed by 72°C for 7 min. The transcriptional levels of each gene were normalized to 16S rRNA.

### Construction of *C. jejuni* mutants and complemented strains

The *dnaK*, *clpB*, *groESL*, *cheY*, *ruvC*, *cmeC*, and *cmeF* knockout mutants were constructed as described previously (61). The *aphA3* (kanamycin resistance) cassette and *cat* (chloramphenicol resistance) cassette were amplified with PCR from pMW10 and pRY112 plasmids, respectively, using primers described in Table S2. Flanking regions of the target genes were also amplified by PCR (Table S2).

Subsequently, the PCR products and pUC19 were digested using BamHI and SalI enzymes, followed by ligation. The resulting plasmids were amplified by inverse PCR and ligated with *aphA3* for constructing *dnaK*, *clpB*, *groESL*, *cheY*, *ruvC*, and *cmeC* mutants or *cat* for the *cmeF* mutant (Table S2). The constructed suicide plasmid was electroporated into *C. jejuni*, and the mutations were confirmed with PCR and sequencing. For the construction of *flaA* and *flaB* mutants, natural transformation was performed as previously described (62). Briefly, the genomic DNAs were extracted from *C. jejuni* 81-176 Δ*flaA*::*cat* (63) and *C. jejuni* 81-176 Δ*flaB*::*cat* mutants in the laboratory collection, digested by SphI and NdeI for Δ*flaA*, and SphI and SalI for Δ*flaB*, respectively. The DNA was spotted directly on the *C. jejuni* cultures grown overnight on MH agar plates and further incubated for 5 h under microaerobic conditions. The bacterial culture was collected and spread on the MH agar plate containing chloramphenicol (12.5 μg/ml). The chloramphenicol-resistant colonies were selected, and the mutations were confirmed by PCR and sequencing using specific primer sets (Table S2). In addition, complemented strains were constructed using the chromosomal integration method (61). The genes (*dnaK*, *clpB*, *groESL*, *ruvC*, *flaA*, *cmeC*, and *cmeF*) were amplified with PCR using primers listed in Table S2, and digested by XbaI or NotI and ligated with the pFMBcomCM plasmid (64) (for *dnaK*, *clpB*, *groESL*, *ruvC*, and *cmeC*), or pFMBcomC (63) (for *flaA*, *cmeF*, and *fur*). The complementation plasmids were introduced to the corresponding mutant strains by electroporation. The complementation into the bacterial chromosome was confirmed with PCR and sequencing. The Δ*fur* mutant strain was previously constructed (65).

### Cross-section transmission electron microscopy (TEM)

*C. jejuni* cells were treated with antibiotics for 24 h, as described above. After harvest, *C. jejuni* cells were washed with ice-cold PBS three times and fixed with Karnovsky’s fixative solution overnight at 4°C. The pellets were washed with 0.05 M sodium cacodylate buffer three times, following post-fixation with 1% osmium tetroxide in the same buffer at room temperature for 1 h. After washing with distilled water three times, en bloc staining was performed with 0.5% uranyl acetate overnight at 4°C. Next day, the samples were washed with distilled water three times, and dehydrated in a series of ethanol gradients (30, 50, 70, 80, 90, and 100%) for 20 min in each step while slowly rotating.

The final 100% ethanol step was repeated three times. Finally, cells were incubated in a 1:1 mixture of Spurr’s resin (66) and ethanol for 90 min at room temperature while slowly rotating and subsequently left in a mixture of 2:1 Spurr’s resin and ethanol at room temperature for 90 min while slowly rotating. The cells were placed in 100% Spurr’s resin and incubated overnight while slowly rotating. The next day, samples were embedded in fresh 100% epoxy resin for 3 h and replaced with fresh 100% epoxy resin. The resin was polymerized for 2 days in an oven at 70°C. The samples cut with an ultramicrotome UC7 (Leica, Wetzlar, Germany) were placed on copper grids and double-stained with 2% uranyl acetate and 3% lead citrate. The sections were observed on a JEM-1010 TEM (JEOL, Tokyo, Japan) operated at 80 kV.

### Confocal fluorescence microscopy

*C. jejuni* cells were treated with antibiotics for 24 h, as described above. *C. jejuni* cells were then washed with ice-cold PBS three times. The pellets were resuspended in 1:500 diluted Proteostat dye (Enzo Life Sciences, NY, USA) and incubated for 20 min in the dark at RT. The cells were simultaneously incubated with SYTO9 (Invitrogen, MA, USA) for 15 min. Then, the sample was washed with PBS. The cells were fixed with 4% paraformaldehyde for 30 min at RT. After washing with PBS, the pellets were resuspended with PBS. 5 μl of each sample was placed on the slides. Confocal images of *C. jejuni* cells were captured using a laser scanning confocal microscope SP8X (Leica, Wetzlar, Germany) using a 488 nm argon laser and a 580 nm emission filter. Images were digitally captured and analyzed with LAS X Software (Leica, Wetzlar, Germany).

### Measurement of intracellular iron levels

The intracellular iron level was measured as previously described (67). Briefly, *C. jejuni* cells were treated with antibiotics for 24 h, as described above. *C. jejuni* cells were then washed with ice-cold PBS three times. After resuspending the pellet with PBS, the cells were disrupted by sonication. Samples were mixed with an iron detection reagent (6.5 mM ferrozine, 6.5 mM neocuproine, 2.5 M ammonium acetate, and 1 M ascorbic acid) and incubated at RT for 30 min. The intracellular iron level was calculated by comparing it with a standard curve obtained with a dilution of 1 mM FeCl3 (Sigma-Aldrich, MO, USA) solution. The absorbance was measured at 550 nm using a SpectraMax i3 platform (Molecular Devices, CA, USA). The intracellular iron level was normalized with the total protein concentrations, which were determined by the Bradford assay (Bio-Rad, CA, USA).

## Data availability

The RNA-seq data have been deposited in the NCBI in the GenBank sequence read archive (SRA) under accession number PRJNA1063286.

## ACKNOWLEDGMENTS

This study was supported by funding from MnDRIVE (Minnesota’s Discovery, Research, and InnoVation Economy) to B.J and the Basic Science Research Program through the National Research Foundation of Korea (NRF) from the Ministry of Education (NRF- 2022R1A6A1A03055869 and NRF-2021R1I1A1A01050990) to J.K.

We declare no competing interest.

B.J. conceptualized the study. E.C., J.K., and J.I.H performed the experiments. B.J. and S.R. supervised the experiments. E.C., J.K., B.J. and S.R. analyzed the results. E.C., J.K., B.J. wrote the manuscript. E.C., and J.K. prepared figures.

