## Supplemental Data for "Pleiotropic cellular responses underlying antibiotic tolerance in *Campylobacter jejuni*"

### Supplemental Materials

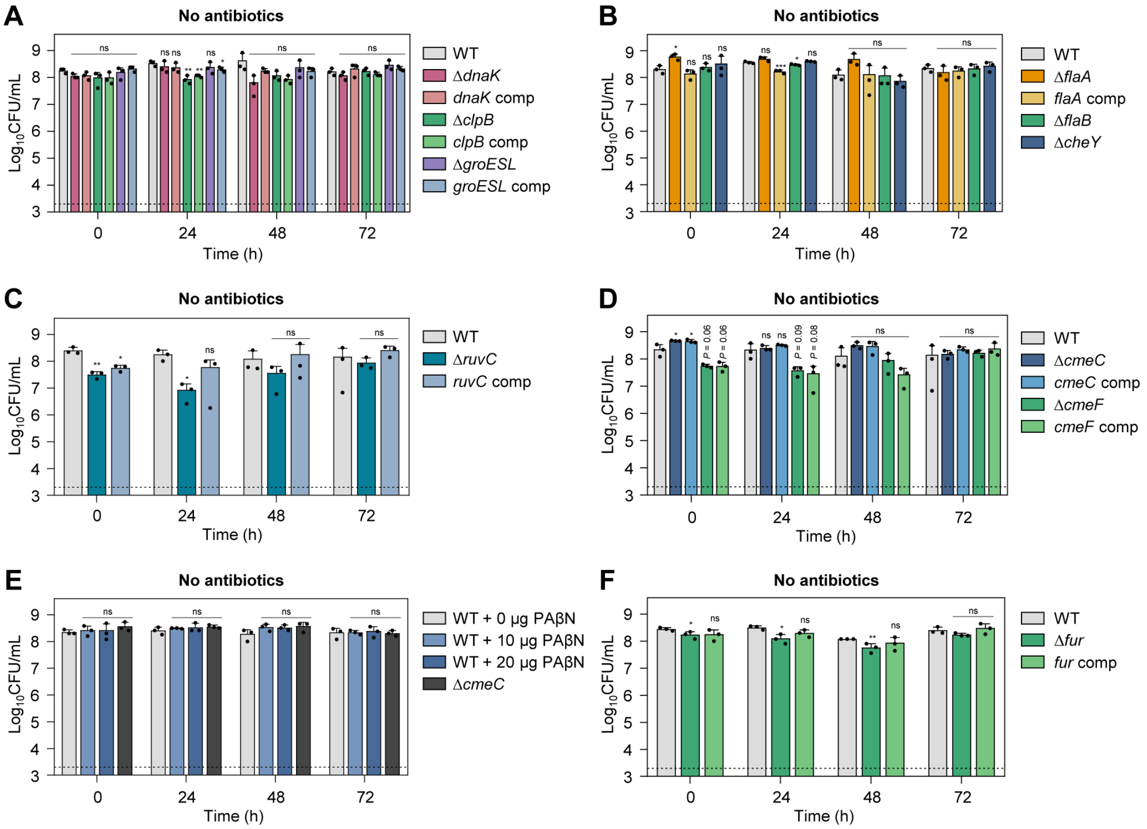

**Fig. S1.** Bacterial survival of the untreated control (No antibiotics) in time-kill assays. Error bars represent the standard deviations of three independent experiments. The data were statistically analyzed by the Student's *t* test in comparison with WT; *, $P < 0.05$; **, $P < 0.01$; ns, not significant; *dnaK* comp, *dnaK*-complemented strain; *clpB* comp, *clpB*-complemented strain; *groESL* comp, *groESL*-complemented strain; *flaA* comp, *flaA*-complemented strain; *ruvC* comp, *ruvC*-complemented strain; *cmeC* comp, *cmeC*-complemented strain; *cmeF* comp, *cmeF*-complemented strain; *fur* comp, *fur*-complemented strain; PAβN, phenylalanine-arginine β-naphthylamide.

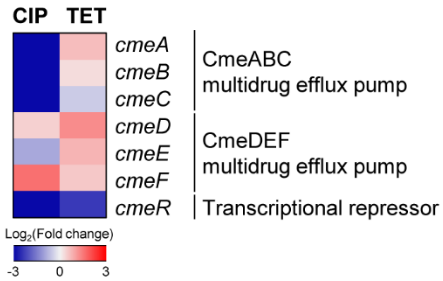

**Fig. S2.** Heat maps of the transcriptional levels of drug efflux pump genes after exposure to 100x MICs of ciprofloxacin (CIP; 6.3 μg/ml) or tetracycline (TET; 3.1 μg/ml). The heat maps were constructed with Gitools.

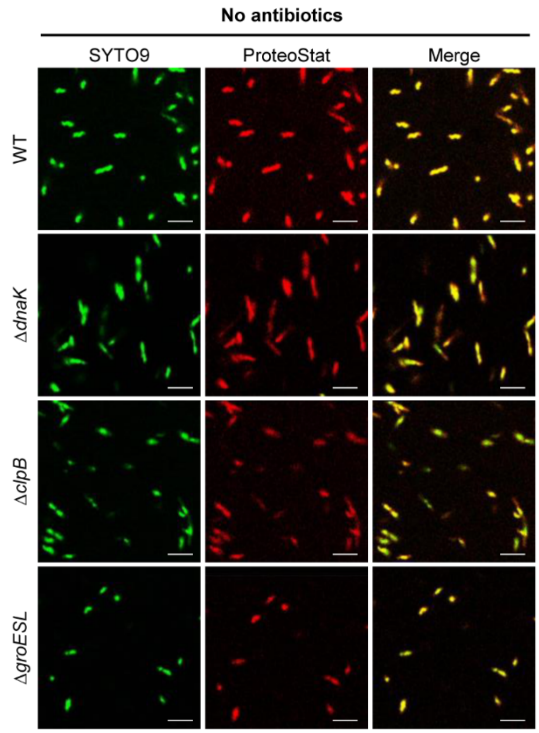

**Fig. S3.** Confocal fluorescence microscopic images of the untreated control (No antibiotics). Live cells were stained with SYTO9 (green), and protein aggregates with ProteoStat® reagent (red). The merged images are shown in yellow. The scale bar represents 5 μm.

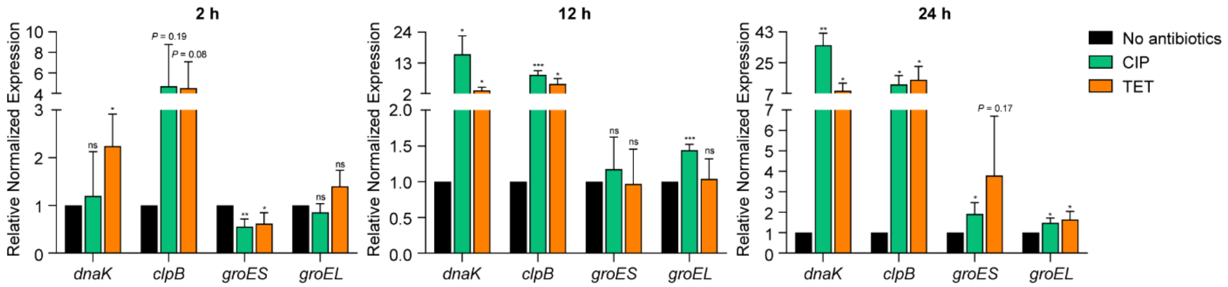

**Fig. S4.** The transcriptional levels of genes associated with chaperones after exposure to 100x MICs of ciprofloxacin (CIP; 6.3 μg/ml) or tetracycline (TET; 3.1 μg/ml). At each time point (2, 12, and 24 h), RNA was extracted and analyzed by qRT-PCR. The expression values were normalized with the untreated control (No antibiotics). Error bars represent the standard deviations of three independent experiments. The data were statistically analyzed by Student's *t* test in comparison with the untreated control (No antibiotics); *, $P < 0.05$; **, $P < 0.01$; ***, $P < 0.001$; ns, not significant.

**Table S1. The minimum inhibition concentrations (MICs) of *Campylobacter* strains**

| Strain | MIC (μg/mL) | |
|---|---|---|
| | **Ciprofloxacin** | **Tetracycline** |
| WT | 0.063 | 0.031 |
| Δ*dnaK* | 0.063 | 0.031 |
| *dnaK* comp | 0.063 | 0.031 |
| Δ*clpB* | 0.063 | 0.031 |
| *clpB* comp | 0.063 | 0.031 |
| Δ*groESL* | 0.063 | 0.031 |
| *groESL* comp | 0.125 | 0.031 |
| Δ*flaA* | 0.063 | 0.031 |
| *flaA* comp | 0.063 | 0.031 |
| Δ*flaB* | 0.063 | 0.031 |
| Δ*cheY* | 0.063 | 0.031 |
| Δ*ruvC* | **0.015** | 0.015 |
| *ruvC* comp | 0.063 | 0.031 |
| Δ*cmeC* | **0.015** | 0.015 |
| *cmeC* comp | 0.031 | 0.015 |
| Δ*cmeF* | 0.063 | 0.031 |
| *cmeF* comp | 0.063 | 0.031 |
| Δ*fur* | 0.063 | 0.063 |
| *fur* comp | 0.063 | 0.031 |

*Changes of at least fourfold are indicated in bold.

**Table S2. Primers used in this study**

| Primer | Sequence (5'-3') | Purpose | Reference |
| --- | --- | --- | --- |
| **Amplifying antibiotic cassette** | | | |
| Kan-F | CTAGCGATGAAGTGCGTAAG | For amplification of *aphA3* (kanamycin resistance) cassette | (1) |
| Kan-R | CGGCTCCGTCGATACTATG | | |
| cat-phospho-F | TGCTCGGCGGTGTTCCTTT | For amplification of *cat* (chloramphenicol resistance) cassette | This study |
| cat-phospho-R | GCGCCCTTTAGTTCCTAAGG | | |
| **Construction of mutant strains** | | | |
| dnaK-SalI-F | AAAGTCGACGCTATAGAAGCAATGAAGAAAGAG | For amplification of *dnaK* with flanking region | This study |
| dnaK-BamHI-R | AAAGGATCCCCATCTCCCCTACTTGAACTG | | |
| dnaK-inv-F | GCGAAGTTTCTCATAAGTTAGCC | For inverse PCR amplification of *dnaK*-cloned pUC19 | |
| dnaK-inv-R | CGTTCATACACAGCAACACAAGA | | |
| clpB-SalI-F | TTTGTCGACGCTTTATCTGCTGGTGTTCCTT | For amplification of *clpB* with flanking region | |
| clpB-BamHI-R | AAAGGATCCGTATGCGTCAACCTGGAAGCT | | |
| clpB-inv-F | CGATATGATACTTGCTGATGAACTT | For inverse PCR amplification of *clpB*-cloned pUC19 | |
| clpB-inv-R | GAATGTATAGCTAAAGATGCTGCA | | |
| groESL-SalI-F | TTTGTCGACGCAATAGGGCTGTAATTATACATTC | For amplification of *groESL* with flanking region | |
| groESL-BamHI-R | AAAGGATCCGCTAGCAATGAAGAAGTAAATACTA | | |
| groESL-inv-F | CGATCCTGTTAAAGTAGAAAGAGTA | For inverse PCR amplification of *groESL*-cloned pUC19 | |
| groESL-inv-R | CTAGAACACGCTTTCCTAAAGG | | |
| cheY-SalI-F | AATGTCGACTGTAGCAAAGAATGAGAAGGTTG | For amplification of *cheY* with flanking region | |
| cheY-BamHI-R | TATGGATCCCTTTCAATAAGCTTTTTTAACTCAGAA | | |
| cheY-inv-F | TAGAACTGTCATCAACAACTAACAA | For inverse PCR amplification of *cheY*-cloned pUC19 | |
| cheY-inv-R | GAAGTGGAGAAGGTGCAGCT | | |

| | | |
|---|---|---|
| ruvC-SalI-F | TTTGTCGACGCATGTTGGGTGGATTGTCTG | For amplification of *ruvC* with flanking region |
| ruvC-BamHI-R | AAAGGATCCCCCCAAAAAGCAGCAGTTTGATT | |
| ruvC-inv-F | CCACAATTTCTCGAACCTGGATC | For inverse PCR amplification of *ruvC*-cloned pUC19 |
| ruvC-inv-R | GGCTTTAACCCATGCAGCAAAT | |
| cmeC-SalI-F | AAAGTCGACTTCAAAACAAAAGCGGAAAAAGCTA | For amplification of *cmeC* with flanking region |
| cmeC-BamHI-R | AATGGATCCGGATAGGTTGTGATTTGTTGCG | |
| cmeC-inv-F | ATTTGAGCAAAGTGAAGATACGAG | For inverse PCR amplification of *cmeC*-cloned pUC19 |
| cmeC-inv-R | CTGAAATCAAAAGAGTAAAACTTGCT | |
| cmeF-SalI-F | ATTGTCGACATGCAAGATTTAGCGAGAGAAAAAAT | For amplification of *cmeF* with flanking region |
| cmeF-BamHI-R | AATGGATCCGTGTTGATTTGATTGATATTTTAGAGG | |
| cmeF-inv-F | CGGTAATAGGTCGGTTTATAGC | For inverse PCR amplification of *cmeF*-cloned pUC19 |
| cmeF-inv-R | CTCGTAGTACCTGCACTTTTTAAA | |

**Confirmation of the mutant strains constructed by natural formation**

| | | | |
|---|---|---|---|
| flaA-con-F | CGATATAGCATTTAACAAGTTCATG | For confirmation of *flaA* gene deletion | This study |
| flaA-con-R | GCAGCTTTTGTAAACTACTGTAG | | |
| flaB-con-F | CAAATCCAAGCCTAGTTTAGAACTA | For confirmation of *flaB* gene deletion | |
| flaB-con-R | CGCTATTTTACCTTTGCTAGACT | | |

**Construction of the complemented strains**

| | | | |
|---|---|---|---|
| dnaK-comp-NotI-F | AAAGCGGCCGCGAGTGGCTCTTATCAAAGATGAAA | For genetic complementation of *dnaK* | This study |
| dnaK-comp-NotI-R | TTTGCGGCCGCGTTTGCTTACACTAAAATGAATTGTCT | | |
| clpB-comp-NotI-F | AAAGCGGCCGCGCAATCGTCTAAGTAGAACCATAG | For genetic complementation of *clpB* | |
| clpB-comp-NotI-R | TTTGCGGCCGCGGAGAAAGTGCTTATTATACCAC | | |
| groESL-comp-NotI-F | AAAGCGGCCGCGCACAACAACAAAAGCTACAATG | For genetic complementation of *groESL* | |
| groESL-comp-NotI-R | TTTGCGGCCGCGGAGGATTTGGTATAGGGCTTT | | |
| ruvC-comp-XbaI-F | ATATCTAGAGTTTACAAGCAAGTTCTGTTCCAA | For genetic complementation of *ruvC* | |
| ruvC-comp-XbaI-R | ATATCTAGATTTTCCAAGTCCTGTAGGTCCA | | |

| | | | |
|---|---|---|---|
| flaA-comp-XbaI-F | AATTCTAGAAAACTTCATATACAAGATAAAACGCAT | For genetic complementation of *flaA* | |
| flaA-comp-XbaI-R | TATTCTAGAGATTAAAGCAAAAAGTGTTCCAAGT | | |
| cheY-comp-XbaI-F | TTTTCTAGATCAGTTTAGTCGTTTGGTATATTTTTG | For genetic complementation of *cheY* | |
| cheY-comp-XbaI-R | ATATCTAGACAAGTGCCCATGAAAACTCTTC | | |
| cmeC-comp-XbaI-F | TATTCTAGACTTTAGCGATATTCTTTGTGCCTT | For genetic complementation of *cmeC* | |
| cmeC-comp-XbaI-R | TATTCTAGAGGCTTATGAAATTACAGATGCAGA | | |
| cmeF-comp-XbaI-F | ATATCTAGAGCTGAAGTGCAAACCACAAATC | For genetic complementation of *cmeF* | |
| cmeF-comp-XbaI-R | TAATCTAGAGAAAAAATACAAATCGCCTATGATAAC | | |
| fur-comp-XbaI-F | GTGGCCTAGGTTTTTAGATCG | For genetic complementation of *fur* | |
| fur-comp-XbaI-R | AAATCTAGAGTCCTGCAACAACAGCATTGA | | |
| **qRT-PCR** | | | |
| 16s-RT-F | ATAAGCACCGGCTAACTCCG | For targeting 16S rRNA | (2) |
| 16s-RT-R | TTCCATCTGCCTCTCCCTCA | | |
| dnaK-RT-F | GGCGAGGTTTTAGTAGGCGA | For targeting *dnaK* | This study |
| dnaK-RT-R | GCGATTTCTATTGCGCACGC | | |
| clpB-RT-F | AGAGCGGGACGAAAAATGGA | For targeting *clpB* | |
| clpB-RT-R | ACGCCTGGTTCACCTAAAAGT | | |
| groES-RT-F | TCAACCTTTAGGAAAGCGTGT | For targeting *groES* | |
| groES-RT-R | TCTGTTCCACCGTATTTAGCA | | |
| groEL-RT-F | AACTATGGGGCCAAGAGGAC | For targeting *groEL* | |
| groEL-RT-R | TGTTCCATCGCCTGCTTGAT | | |

*Underlining indicates the enzyme recognition sites.
